# Erythrocyte aging links hypocalcemia to impaired flow dynamics and oxygen transport through volume dysregulation

**DOI:** 10.64898/2026.10.26.762074

**Authors:** Jiaqi Lim, Justin Kok Soon Tan, Yijia Yuan, Elaine Shi An Chen, Yujin Song, Sim Leng Tien, Hwa Liang Leo, Sangho Kim

**Affiliations:** Department of Biomedical Engineering, National University of Singapore, Singapore 119077; NUS Graduate School – Integrative Sciences and Engineering Program, National University of Singapore, Singapore 119077; N.1 Institute for Health, National University of Singapore, Singapore 119077; Department of Hematology, Singapore General Hospital, Singapore 169608

## Abstract

Red blood cells (RBCs) maintain their corpuscular volume through mechanotransducive ion-mediated processes in response to circulatory shear stress. However, with age, RBCs accumulate oxidative damage and undergo alterations in protein function, glucose metabolism and ion transport, raising fundamental questions about how volume homeostasis is preserved under mechanical load. Here, we investigated the differential responses of younger (YRBCs) and older RBCs (ORBCs) to fluid shear stress, with a focus on calcium-dependent regulation. We found that ORBCs exhibited a striking dependence on extracellular calcium to maintain volume stability. In calcium-depleted conditions, only ORBCs underwent pronounced post-shear swelling, with recovery only partially restored by ATP-dependent mechanisms. In contrast, calcium preserved volume homeostasis, identifying calcium-sensitive pathways as key compensatory regulators in metabolically compromised ORBCs. Notably, metabolically deficient YRBCs recapitulated the ORBC swelling phenotype, supporting a shift from ATP to calcium-dependent regulation with age. More importantly, shear-induced swelling impaired flow behaviour and oxygen delivery in ORBCs, linking defective volume control to microcirculatory dysfunction. Together, these findings potentially established calcium-dependent volume regulation as a critical adaptive mechanism in ageing RBCs and reveal a previously unrecognized hematological link to vascular dysfunction in hypocalcemia.

## Introduction

Red blood cells (RBCs) are primarily responsible for the transport of oxygen (O_2_) from the lungs to tissue via the systemic circulation. One important biophysical property is their corpuscular volume, maintained through a robust system of ion channels. RBCs have a unique biconcave discoid shape^1^ with a mean corpuscular volume (MCV) of ∼90 fL and a mean surface area of ∼135 μm^2^, corresponding to the surface area to volume ratio of ∼60%^2^ that is preserved and strictly maintained throughout their lifespan^3,4^. Deviation from this means RBCs becoming swelled or dehydrated, impairing their deformability and other biophysical traits^5,6^. As RBCs circulate in the body, they are constantly exposed to a wide range of shear stresses in the cardiovasculature^7–9^. The shear stress-induced deformation of the cell membrane is associated with the activation of the mechanosensitive, nonspecific cation channel, Piezo1^10,11^. The opening of these channels increases the permeability of the RBC membrane to an influx of monovalent cations (sodium, Na^+^ and potassium, K^+^) and calcium (Ca^2+^), resulting in a fluctuation of cytosolic cationic composition^12,13^. Since cellular hydration is predominantly controlled by movement of monovalent cations^14^, the exposure to shear stress may significantly alter RBC volume. On the other hand, while Ca^2+^ does not directly influence volume change in RBCs, influx of Ca^2+^ ions into the RBC triggers the activation of the Ca^2+^-sensitive, K^+^-selective Gardos channel. The secondary activation of the Gardos channel mediated by Ca^2+^ results in an outward displacement of K^+^ and water from the RBC, effectively dehydrating the cell^15,16^. RBCs maintain a basal cytosolic Ca^2+^ level through their low membrane permeability to Ca^2+^ and plasma membrane Ca^2+^ ATPases (PMCA) responsible for removing excess levels of Ca^2+^ within the RBCs^17^ to prevent the excessive loss of water.

As RBCs age, they accumulate oxidative damage as an inevitable consequence of their exposure to endogenous and exogenous reactive O_2_ species^18^. Despite the extensive antioxidant faculties that the RBCs possess^19,20^, the vulnerability to oxidative stresses increases with age caused by the limited capacity to replace damaged cellular components. One primary age-related change would be the loss of glucose metabolism. The lack of critical enzymes necessary for glycolysis results in a significant reduction in adenosine triphosphate (ATP) availability. The reduction of ATP production by up to 40%^21^ in older RBCs (ORBCs) is likely to result in a decline of ORBC volume control. Primarily, membrane ion pumps such as Na^+^ K^+^ Pump (ATP1) and PMCA which rely on ATP as an energy source will have impeded function. This is aggravated by a reduction of up to 70% in the number of ATP1 sites in ORBCs^22^ as compared to that in younger RBCs (YRBCs). Collectively, the decreased number of ATP1 and gradual attenuation in the activity of the remaining ATP1 may likely impair the cell’s ability to regulate its volume. Interestingly, while the number of ATP1 decreases drastically with age, the number of Gardos channels seems to be maintained on the RBC membrane^23,24^, suggesting the potential role of Ca^2+^ as a primary means of volume control in ORBCs to compensate for the inevitable decline in ATP1 activity.

Hypocalcemia can occur for several reasons and is often characterized by insufficient levels of parathyroid hormone or vitamin D^25–28^. This significant depletion of Ca^2+^ in blood is often associated with chronic kidney disease (CKD)^29^. Globally, CKD is a prevalent condition that affects 1 in 10 people^30^. Furthermore, there is a strong correlation between CKD and cardiovascular diseases (CVD)^31^, suggesting a potential role that low Ca^2+^ has in CVD pathogenesis. The relationship between hypocalcemia and CVD is complex and indirect^32,33^, encompassing direct impairments in cardiac function^34^ and cumulative effects which contribute to CVD pathogenesis. However, existing studies have neglected the effects of low Ca^2+^ on the biophysical properties and function of RBCs. Specifically, there has been little information on the possibility of hypocalcemia impacting hemodynamics and O_2_ transport. With the sheer abundance of RBCs in circulation, the changes in RBC properties caused by impaired volume regulation might also be a contributing factor to CKD and CVD aetiology. Hence, our study puts forth a hematological perspective in understanding the disease conditions occurring in hypocalcemic patients.

Thus far, there has been no direct evidence on the extent of RBC volume change in hypocalcemic conditions or on how Ca^2+^ presence might mitigate this effect. Our findings reveal a hitherto unknown distinction in responses to shear-induced swelling between YRBCs and ORBCs when sheared in the absence of Ca^2+^. Specifically, only ORBCs exhibited a significant increase in size after shear treatment. The enlarged ORBCs gradually shrank over time, albeit not being able to fully recover their initial size. Notably, this volume recovery seemed to be ATP-dependent since metabolic-deficient YRBCs also swelled after shear treatment. Interestingly, no swelling was observed for both subpopulations of RBCs when sheared in the presence of Ca^2+^. We further demonstrated that the swelling of sheared ORBCs significantly altered their flow dynamics and O_2_ diffusion, alluding to possible microcirculation impairment as a consequence of the volume dysregulation in a hypocalcemic environment.

## Results

### Volume regulation is impaired in ORBCs in the absence of Ca^2+^

We used the two-dimensional area data to represent RBC size and this analysis was validated by comparison with three-dimensional mean corpuscular volume (MCV) data. The detailed information has been provided in the methodology section and in **Supplementary Figures. S1 and S2**. As expected, we observed a distinct difference in size between different biologically aged RBCs similar to previous studies^35,36^. Both YRBCs and ORBCs followed a normal distribution with YRBCs found to be ∼11% larger (YRBCs: 49.4 ± 0.51 µm^2^ vs ORBCs: 44.5 ± 0.48 µm^2^, p < 0.001) (**Figure. 1a**). The normality of the respective RBC subpopulations was found to be preserved before and after shear treatment (**Figure.1b**). In the case of YRBCs, no significant change in RBC size was observed when suspended in media with and without Ca^2+^ (with Ca^2+^: 49.3 ± 0.53 µm^2^ vs without Ca^2+^: 49.4 ± 0.51 µm^2^, p > 0.05). There were also no significant changes in the distribution profile and size before shearing (BS) and after shearing (AS) regardless of the suspending media (with Ca^2+^, BS: 49.3 ± 0.53 µm^2^ vs AS: 49.3 ± 0.54 µm^2^, p > 0.05; without Ca^2+^, BS: 49.4 ± 0.51 µm^2^ vs AS: 49.5 ± 0.52 µm^2^, p > 0.05) (**Figure.1c**). The use of different media (with and without Ca^2+^) also showed no impact on the ORBC size (with Ca^2+^: 44.4 ± 0.56 µm^2^ vs without Ca^2+^: 44.5 ± 0.48 µm^2^, p > 0.05). However, while ORBCs showed no significant change in size after being sheared in media with Ca^2+^ (BS: 44.4 ± 0.56 µm^2^ vs AS: 44.4 ± 0.56 µm^2^, p > 0.05) similar to YRBCs, there was an apparent rightward shift in the distribution profile corresponding to a significant increase in the mean RBC size by 3.5 ± 0.08% after shear exposure in the absence of Ca^2+^ (BS: 44.5 ± 0.48 µm^2^ vs AS: 46.0 ± 0.50 µm^2^, p < 0.001) (**Figures.1b and 1d**). As reduction in RBC volume is associated with the efflux of monovalent cations and water, medium K^+^ was selected as an indicator due to its association with the Gardos channel. A significant increase in medium K^+^ levels was found in both subpopulations of RBCs after the shear treatment in both media (with and without Ca^2+^) (p < 0.05). However, in the case of ORBCs sheared in media without Ca^2+^, there was a significantly smaller increase in medium K^+^ (10.0 ± 0.11%, p < 0.05) as compared to the rest of the cases (**Figure. 1e**).

**Figure. 1:**
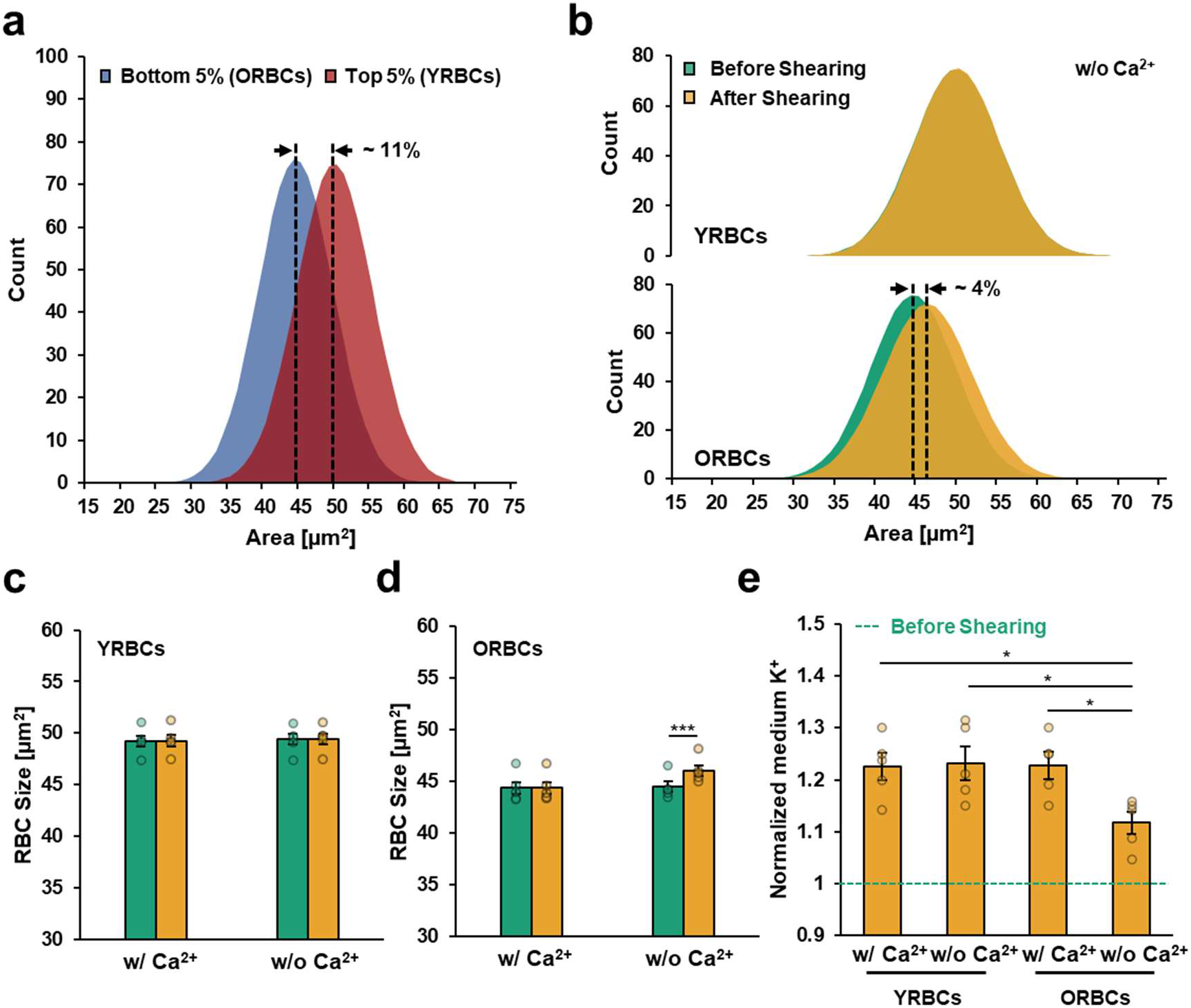
Age-dependent RBC swelling in response to shear stress. **a:** Normal distributions of isolated RBC populations (i.e. YRBCs and ORBCs) showing the significantly larger mean cell size in YRBCs. **b:** Visualization of the shift in size distribution after shear treatment in the absence of Ca^2+^. ORBCs exhibited a significant increase in size after shearing. **c:** Effects of shearing on the size of YRBCs. No significant change in size was observed after shear treatment regardless of the media composition. **d:** Effects of shearing on the size of ORBCs. ORBCs sheared in the absence of Ca^2+^ showed significant swelling after shear treatment. **e:** Comparison of normalized medium K^+^ levels after shear treatment. The green dashed line represents the normalized baseline for each condition before the shear treatment. *P* value in a for size comparison between YRBCs and ORBCs was calculated using unpaired t-test. *P* values in c and d were calculated using two-way repeated measures ANOVA with Tukey’s post-hoc test. *P* value in e was calculated using one-way ANOVA with Tukey’s post hoc test. Data presented as means ± SEM; n = 5 of biologically independent sets of RBCs. \**P* < 0.05 \*\*\**P* < 0.001.

### ATP is essential for effective volume regulation in the absence of Ca^2+^

Fresh RBCs and metabolic-deficient (MD) RBCs were compared to investigate the effects of decreased ATP on shear-induced swelling in the absence of Ca^2+^. Additionally, the size of the RBCs was monitored for 40 min after shear cessation for any potential size-recovery of the swelled RBCs after shear treatment. There was no significant difference in cell deformability between fresh and MD RBCs (**Supplementary Table S1**). A significant reduction in ATP levels was observed in both subpopulations of RBCs with MD YRBCs and MD ORBCs experiencing a 57.6 ± 3.37% (p < 0.01) and 68.5 ± 4.89% (p < 0.01) decrease in ATP, respectively (**Figure. 2a**). This affirmed the metabolic deficiency in the prepared MD RBCs. Fresh YRBCs showed no significant swelling after shear treatment and there was no significant change in their size from baseline across all the measured time interval (p > 0.05). In contrast, the MD YRBCs showed an initial increase in size by 2.4 ± 0.14% followed by a gradual size recovery with time, characterized by an exponential curve (*_P_* = 2.20e^-0.150t^ + 0.120, R^2^ = 0.980). On the other hand, both fresh and MD ORBCs swelled after shearing to a similar degree (fresh ORBCs: 3.8 ± 0.05% vs MD ORBCs: 3.7 ± 0.09%, p > 0.05). However, the recovery behaviour for both cases were very different, following two separate exponential decay functions (fresh ORBCs: *_P_* = 2.11e^-0.210t^ + 1.60, R^2^ = 0.965 vs MD ORBCs: *_P_* = 1.32e^-0.08t^ + 2.34, R^2^ = 0.977) (**Figure. 2b**). As fresh YRBCs did not show any swelling after shear treatment and was able to retain the same size, the recovery potential (*RP*) was assumed to be 100%. MD YRBCs had a reduced RP (92.9 ± 3.05%) but was not significantly different from the fresh YRBCs (p > 0.05). Comparing the fresh YRBCs and ORBCs, the fresh ORBCs were not able to recover fully and had a significantly lower RP of 57.5 ± 1.79% (p < 0.001). The *RP* was further reduced in the MD model (36.3 ± 4.91%), significantly lower than its fresh counterpart (p < 0.01) (**Figure. 2c**). Comparing the recovery rate constant (τ), the τ of fresh ORBCs was significantly higher than fresh YRBCs (ORBCs: 5.22 ± 0.843 vs YRBCs: 0.103 ± 0.0552, p < 0.01), highlighting that ORBCs displayed a slower size recovery. A significant increase in τ was observed in MD YRBCs (fresh: 0.103 ± 0.0552 vs MD: 6.18 ± 1.100, p < 0.001). Similarly, there was a significant increase in τ observed in MD ORBCs (fresh: 5.22 ± 0.843 vs MD: 12.4 ± 1.07, p < 0.001) (**Figure. 2d**).

**Figure. 2:**
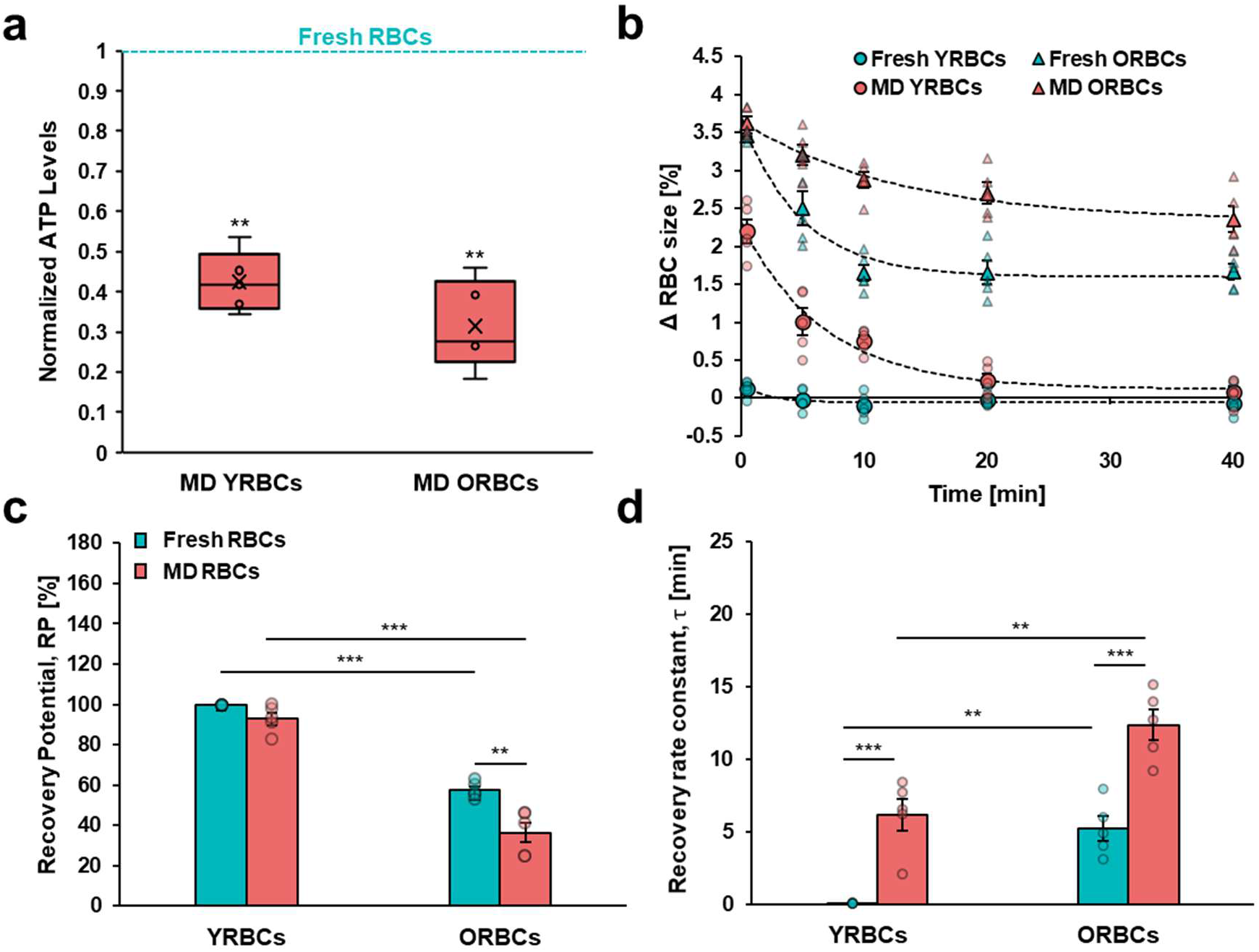
Fresh RBCs vs MD RBCs in size recovery post-shear induced swelling. **a:** Normalized ATP levels in MD RBCs. The turquoise dashed line denotes the normalized baseline, each ATP level in fresh (non-MD) RBCs (*P* values for comparison between fresh and MD RBCs were calculated using paired t-tests). **b:** Recovery profile of RBCs after shear treatment for a 40-min duration. Exponential curves were fitted for the data that showed initial swelling and subsequent recovery. Black dashed lines were used to illustrate the non-linear regression fit (R^2^_MDYRBC_ = 0.980, R^2^_freshORBC_ = 0.965, R^2^_MDORBC_ = 0.977). The recovery rate constant (τ) and recovery potential (*RP*) information was derived from the fitted curves. **c:** Comparison of RP between fresh and MD RBCs of YRBCs and ORBCs. *RP* was shown to be reduced in MD cells regardless of biological age. **d:** Comparison of τ between fresh and MD RBCs of YRBCs and ORBCs. τ was elevated in both MD groups, with the ORBCs consistently showing a higher τ in both fresh and MD groups, representing a slower recovery of cell size post-shear treatment. *P* values in c and d were calculated using two-way repeated measures ANOVA with Tukey’s post-hoc test. Data presented as means ± SEM; n = 5 of biologically independent sets of RBCs. **P < 0.01 ***P < 0.001.

### Gardos effect modulates volume regulation in ORBCs

The average fluorescence intensity obtained from Fluo-4 loaded RBCs were compared before and after shear treatment in both YRBCs and ORBCs to determine the retained intracellular Ca^2+^. Brightfield and the corresponding fluorescent images were captured after shearing to visualize the retention of intracellular Ca^2+^ in YRBCs and ORBCs, respectively (**Figure. 3a**). Both subpopulations of RBCs were found to have an increase in intracellular Ca^2+^ after shear treatment than their respective baseline values (YRBCs: 19.4 ± 6.66% vs ORBCs: 39.0 ± 2.93%). The change in fluorescence intensity after shear treatment was significantly different between YRBCs and ORBCs (p < 0.05) (**Figure. 3b**). ORBCs were used to test for the degree of swelling when sheared in different Ca^2+^ concentrations. Interestingly, there was a strong negative linear correlation between the media Ca^2+^ concentration and the shear induced swelling of ORBCs (y = -1.83x + 3.785, R^2^ = 0.985) (**Figure. 3c**). Our results revealed the significance of Ca^2+^ in the maintenance of RBC volume, especially for ORBCs. More importantly, it should be noted that a Ca^2+^ level of at least 2 mM is necessary for optimal volume regulation. To confirm the contributions of Ca^2+^ in volume regulation, inhibition of the Ca^2+^-activated Gardos channels was carried out with the use of Senicapoc. Our results showed no significant difference in size between control and Gardos inhibited YRBCs (control YRBCs: 0.0206 ± 0.08261% change vs Gardos inhibited YRBCs: 0.0626 ± 0.06640% change, p > 0.05). However, in the case of ORBCs, there was a significant difference between the control and Gardos inhibited samples (control ORBCs: 0.0443 ± 0.06204% change vs Gardos inhibited ORBCs: 3.69 ± 0.070% change, p < 0.001), confirming the dominant role that Ca^2+^ plays in regulating volume in ORBCs (**Figure. 3d**).

**Figure. 3:**
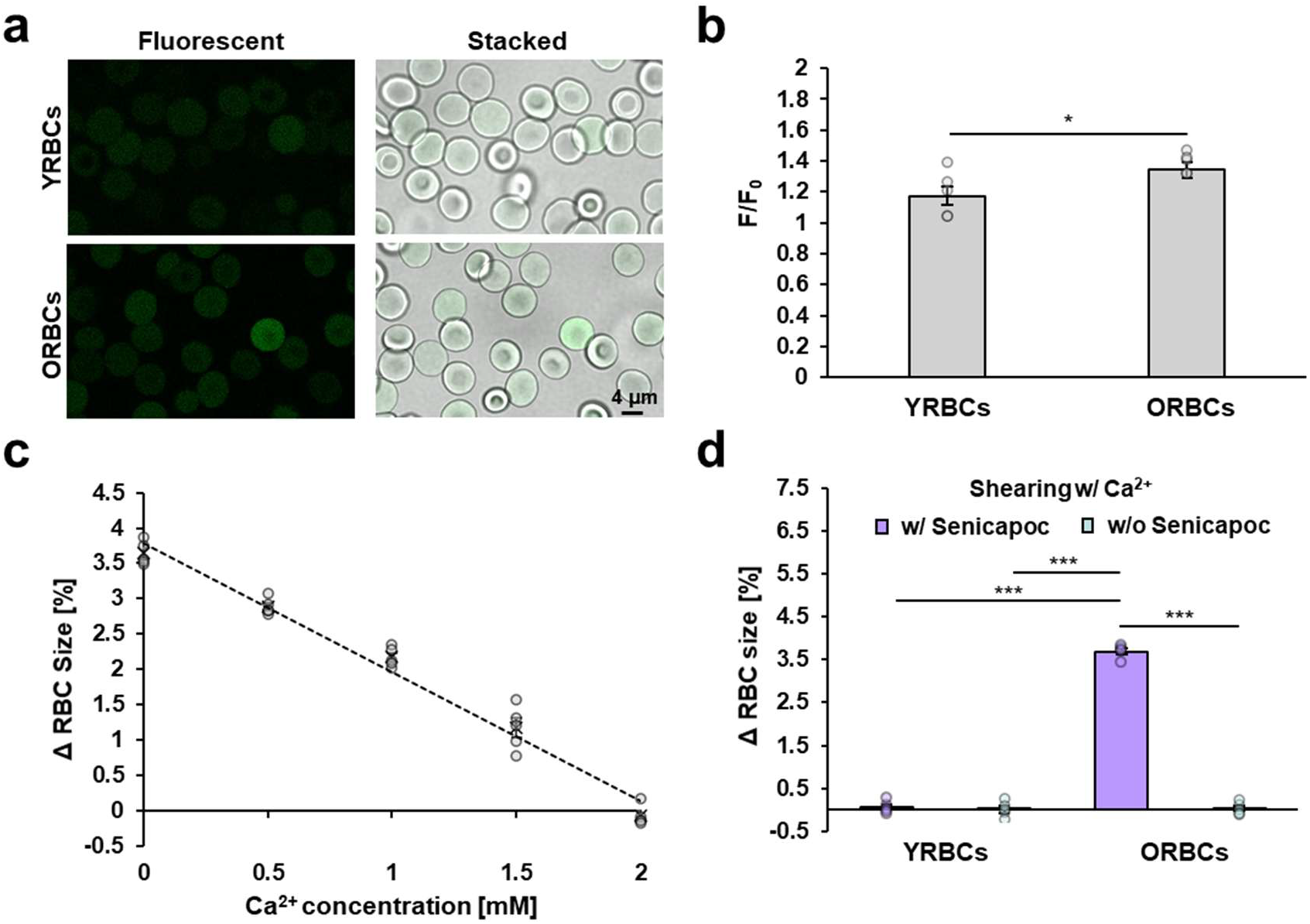
Dependence of different biologically aged RBCs on Ca^2+^ in size recovery post-shear induced swelling. **a:** Qualitative images comparing the Fluo4-AM fluorescence emitted from YRBCs and ORBCs after shear treatment. **b:** Normalized Fluo4-AM fluorescence emitted from YRBCs and ORBCs with ORBCs showing a higher fluorescence after shear treatment (*P* value for comparison of fluorescence after shearing between YRBCs and ORBCs was calculated using unpaired t-test). **c:** Dose-dependent effects of extracellular Ca^2+^ on shear induced ORBC swelling. The dashed line corresponds to the linear regression fit of the data (R^2^ = 0.985) **d:** Effects of Senicapoc on shear induced swelling of YRBCs and ORBCs in Ca^2+^ media. *P* values in d were calculated using two-way repeated measures ANOVA with Tukey’s post-hoc test. Data presented as means ± SEM; n = 5 of biologically independent sets of RBCs. * p < 0.05. *** p < 0.001.

### Shear-induced swelling affects flow dynamics of ORBCs

Microvascular bifurcations are ubiquitous in the arteriolar network and are instrumental to the distribution of blood to the peripheral circulation. At this length scale, RBC biophysics can modulate the flow dynamics and cell partitioning, affecting the overall tissue perfusion efficiency. To investigate the potential effect of the shear-induced swelling on hemodynamics within the arteriolar network *in vitro*, we utilized a microfluidic platform (**Figure. 4a**) to simulate blood flow in an arteriolar bifurcation. A detailed description of the microfluidic channel design has been provided in **Supplementary Figure. S3**. Upstream positions of RBCs in each subpopulation were analyzed up to 3 channel widths (3D) away from the bifurcation point to confirm that the starting positions of RBCs were not significantly different between cases (**Supplementary Figure. S4**).

**Figure. 4:**
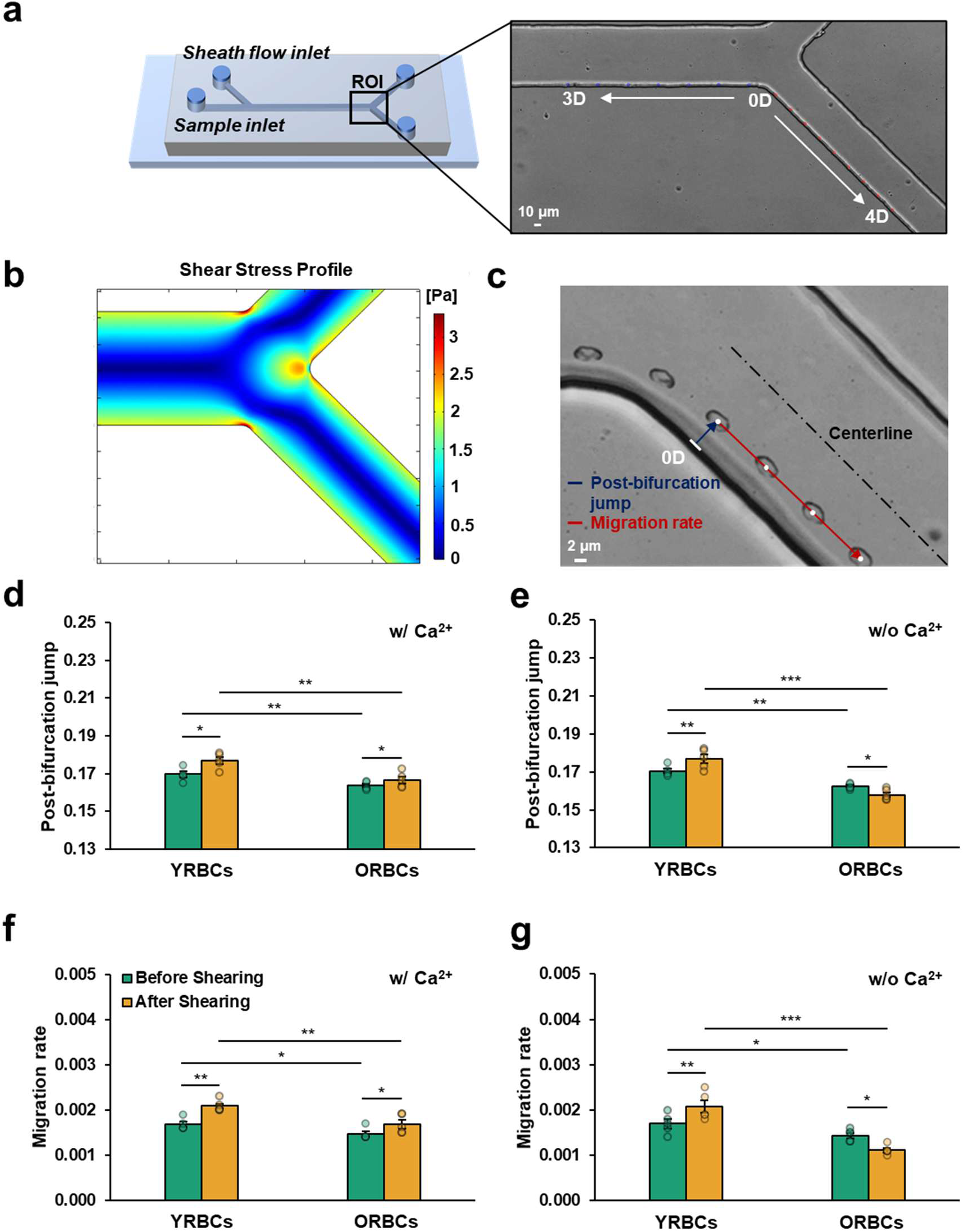
Single cell dynamics quantified with an in vitro microfluidic channel. **a:** Schematic of the bifurcating channel. The sheath flow is responsible for initializing the positions of the RBCs towards the inner wall of the channel prior to the bifurcation. Distance of RBCs away from the channel walls were quantified at every 0.5D for 3D upstream and 4D downstream as a normalized distance of the channel width (D: channel width). **b:** Two-dimensional COMSOL simulation with geometry mimicking the bifurcating channel was performed to obtain the shear stress profile at the bifurcation. **c:** Zoomed-in stacked image of a single RBC trajectory past the bifurcation zone. The post-bifurcation jump (blue arrow) was obtained by normalizing downstream particle position with upstream initialized position. The migration rate (red arrow) was determined from the gradient of the linear fit obtained from the cell movement downstream up to 4D. **d:** Migration rate of YRBCs and ORBCs before and after shearing in the presence of Ca^2+^. **e:** Migration rate of YRBCs and ORBCs before and after shearing in the absence of Ca^2+^. **f:** Post-bifurcation of YRBCs and ORBCs before and after shearing in the presence of Ca^2+^. **g:** Post-bifurcation of YRBCs and ORBCs before and after shearing in the absence of Ca^2+^. *P* values in d-g were calculated using two-way repeated measures ANOVA with Tukey’s post-hoc test. Data presented as means ± SEM; n = 5 of biologically independent sets of RBCs. *P < 0.05 **P < 0.01 ***P< 0.001.

The near wall shear stress profile for the bifurcation model was obtained from computational fluid dynamics simulations at the bifurcation region (**Figure. 4b**). We found that the RBCs were exposed to a shear stress of ∼1.5 Pa in our *in vitro* experiments. RBC deformability index (DI) at 1.5 Pa before and after shearing in both media (with and without Ca^2+^) were measured (**Supplementary Table S2**) for comparison with RBC post-bifurcation dynamics. In all cases, the deformability of RBCs was significantly higher in the treated samples (p < 0.05). This is with the exception of the case when ORBCs were sheared without Ca^2+^ where their deformability was significantly lower (p < 0.05). The trajectories of RBCs post-bifurcation across all cases were recorded. The downstream lateral positions of the RBCs were then determined at every 0.5D and normalized with the channel width (D). For each case, the downstream positions of at least 100 single RBCs across 4D from the bifurcation point were analyzed for comparison. For this experiment, two parameters were derived from the linear fit of the RBC trajectories downstream of the bifurcation (y = *m*x + *j*). As illustrated in **Figure. 4c**, the displacement of the RBC away from the wall at 0D of the daughter channel is represented as post-bifurcation jump (*j*) while the rate at which the RBC migrates towards the centreline of the flow as migration rate (*m*). The application of shear treatment was observed to increase both *j* and *m* of RBCs regardless of cell age. This is with exception for the case of ORBCs sheared in the absence of Ca^2+^. In this case, shear treatment significantly reduced both *j* and *m* (**Figures. 4e and 4g**). It is of note that flowing the respective RBC subpopulations in either medium showed no significant difference in *j* or *m* (p > 0.05). With Ca^2+^ before shearing, *j* and *m* were significantly higher in YRBCs than the ORBCs (*j* - YRBCs: 0.172 ± 0.0010 vs ORBCs: 0.163 ± 0.0011, p < 0.01; m - YRBCs: 0.00178 ± 0.000080 vs ORBCs: 0.00146 ± 0.000060, p < 0.05). Similarly, without Ca^2+^ before shearing, *j* and *m* were also significantly higher in YRBCs than the ORBCs (*j* - YRBCs: 0.172 ± 0.0012 vs ORBCs: 0.163 ± 0.0009, p < 0.01; *m* - YRBCs: 0.00180 ± 0.000130 vs ORBCs: 0.00146 ± 0.000051, p < 0.05). After shearing, in the absence of Ca^2+^, sheared YRBCs showed a significant increase in *j* (BS: 0.172 ± 0.0012 vs AS: 0.180 ± 0.0015, p < 0.01) and *m* (BS: 0.00180 ± 0.000130 vs AS: 0.00224 ± 0.000121, p < 0.01). On the contrary, ORBCs showed a significant decrease in both *j* (BS: 0.163 ± 0.0009 vs AS: 0.158 ± 0.0008, p < 0.05) and *m* (BS: 0.00146 ± 0.000051 vs AS: 0.00116 ± 0.000040, p < 0.05). Intriguingly, when sheared with Ca^2+^, ORBCs displayed a reverse trend and exhibited a significant increase in both *j* (BS: 0.163 ± 0.0011 vs AS: 0.168 ± 0.0015, p < 0.05) and *m* (BS: 0.00146 ± 0.000060 vs AS: 0.00178 ± 0.000097, p < 0.05). On the contrary, YRBCs showed the same increasing trend in both *j* (BS: 0.172 ± 0.00098 vs AS: 0.178 ± 0.0010, p < 0.05) and *m* (BS: 0.00178 ± 0.000080 vs AS: 0.00224 ± 0.000081, p < 0.01).

### Shear-induced swelling impairs O_2_ transport of ORBCs

The primary role of RBCs in the circulation is to deliver O_2_. To evaluate the overall efficacy of O_2_ delivery, p50 and O_2_ release rate were quantified. The p50 provides information on the capability of RBCs to carry O_2_ by measuring the affinity of hemoglobin for O_2_. On the other hand, RBC O_2_ release rate quantifies the diffusion speed of O_2_ once dissociated from hemoglobin. In the case of p50, YRBCs was significantly higher than ORBCs regardless of the suspending media (with Ca^2+^ - YRBCs: 27.1 ± 0.51 mmHg vs ORBCs: 24.9 ± 0.37 mmHg, p < 0.05; without Ca^2+^ - YRBCs: 27.1 ± 0.16 mmHg vs ORBCs: 25.3 ± 0.26 mmHg, p < 0.05). Within the same RBC subpopulations, there were no significant differences in p50 when RBCs were suspended in different media for both YRBCs and ORBCs (p > 0.05). Interestingly, the exposure to shear treatment resulted in a significant decrease of p50 for all cases (p < 0.05). It is of note that the regardless of the RBC subpopulation and suspending media, the sheared RBCs in all cases showed a similar reduction of p50 to ∼23 mmHg (**Figures. 5a and 5b**). To analyze the RBC O_2_ release rate, an integrated system comprising gas delivery tubes and an electrochemical O_2_ probe was designed and utilized (**Figure. 5c**). The detailed description of the system and determination of O_2_ release rate has been provided in the methodology section. To validate the system’s performance, a comparison was first made between PBS and RBCs suspended in PBS at a hematocrit of 1% (**Figure. 5d**). Both samples were initialized to 283 µmol/L with sterile air prior to the deoxygenation process with nitrogen gas. The deoxygenation process of both cases followed an exponential decay (PBS: *_C_* = 283e^-0.0248t^, R^2^ = 0.998 vs RBCs in PBS: *_C_* = 283e^-0.0201t^, R^2^ = 0.994). In the presence of RBCs, the deoxygenation process of the suspending media was significantly slower (p < 0.001), evident from the reduced decay constant from the fitted exponential curve. As shown in **Figures. 5e and 5f**, the O_2_ release rate of YRBCs was significantly lower than that of ORBCs in both suspending media (with Ca^2+^ - YRBCs: 0.0179 ± 0.00011 s^-1^ vs ORBCs: 0.0201 ± 0.00022 s^-1^, p < 0.001; without Ca^2+^ - YRBCs: 0.0179 ± 0.00017 s^-1^ vs ORBCs: 0.0201 ± 0.00013 s^-1^, p < 0.01). This indicates that ORBCs are capable of releasing O_2_ faster than YRBCs and the presence of Ca^2+^ has no effect on the basal O_2_ release rate. While the shear treatment did not result in a significant effect on the O_2_ release rate across majority of the cases tested (p > 0.05), ORBCs sheared in the absence of Ca^2+^ showed a significantly lower O_2_ release rate (BS: 0.0201 ± 0.00013 s^-1^ vs AS: 0.0190 ± 0.00028 s^-1^, p < 0.05).

**Figure. 5:**
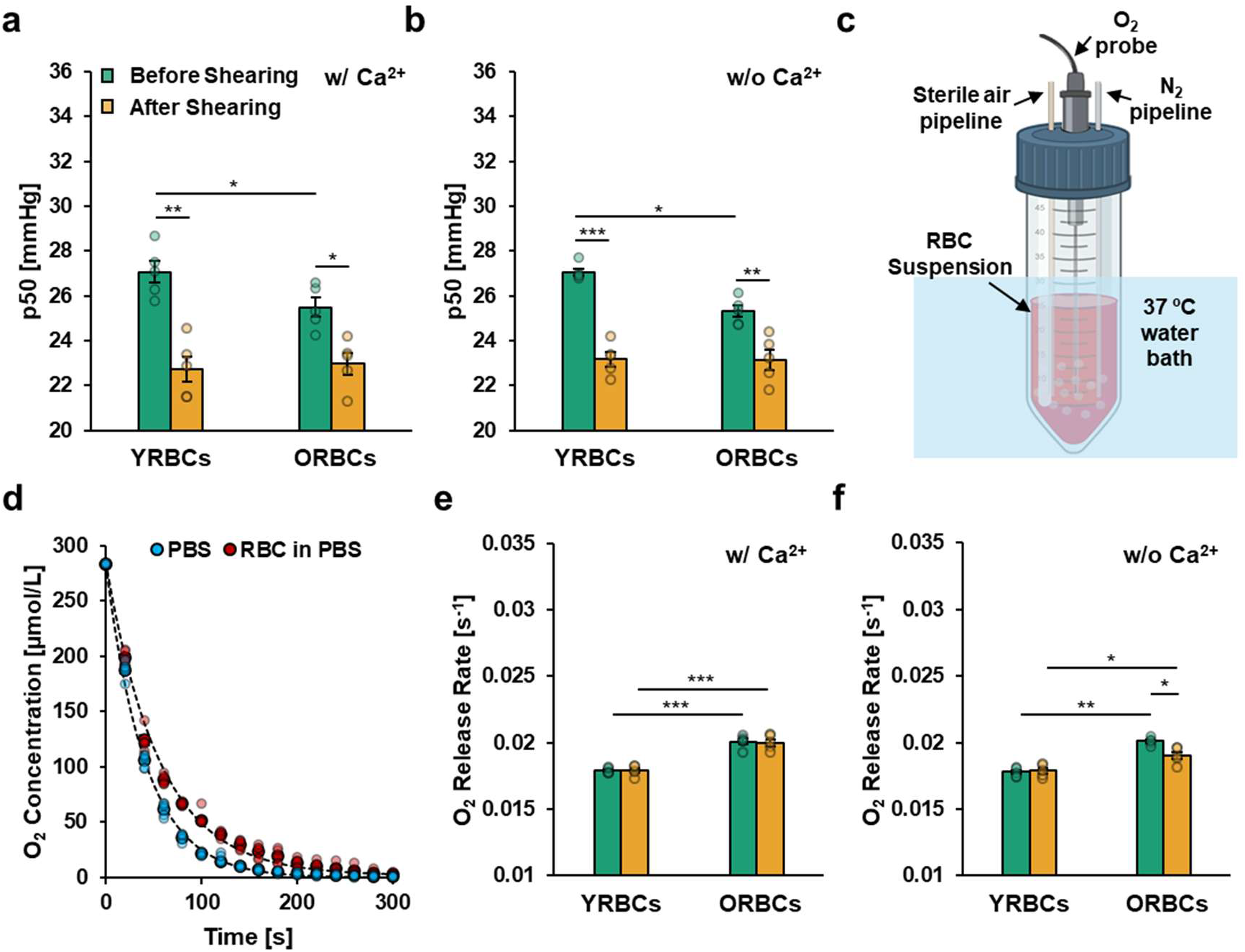
p50 and O_2_ release rate in relation to change of RBC age and size. **a:** p50 of YRBCs and ORBCs before and after shearing in the presence of Ca^2+^**. b:** p50 of YRBCs and ORBCs before and after shearing in the absence of Ca^2+^**. c:** Schematic diagram showing the in-house set up for determination of RBC O_2_ release rate. **d:** Comparison of deoxygenation rates in the presence and absence of RBCs. The presence of RBCs in the solution resulted in a slower deoxygenation rate of the suspending media. The deoxygenation process can be described with a non-linear regression fit as illustrated with black dashed lines (R^2^_PBS_ = 0.998, R^2^_RBC_ = 0.994). (*P* value for comparison of O_2_ release rate between PBS and RBC suspension was calculated using unpaired t-test) **e:** O_2_ release rate of YRBCs and ORBCs before and after shearing in the presence of Ca^2+^**. f:** O_2_ release rate of YRBCs and ORBCs before and after shearing in the absence of Ca^2+^. *P* values in a, b, e and f were calculated using two-way repeated measures ANOVA with Tukey’s post-hoc test. Data presented as means ± SEM; n = 5 of biologically independent sets of RBCs. *P < 0.05 **P < 0.01 ***P < 0.001.

## Discussion

In our work, we report a shift in volume regulatory shear response between YRBCs and ORBCs that can be dichotomized into Ca^2+^ independent (ATP-driven active pathways and passive transporters) and Ca^2+^ dependent (Gardos channel) mechanisms. Primarily, we highlighted the importance of Ca^2+^ in volume regulation, which was crucial for maintaining RBC flow dynamics and O_2_ delivery efficiency, especially in ORBCs. Our findings suggest that the need to sustain RBC functionality has likely driven the evolution of an energy-efficient compensation strategy in volume regulation (**Figure. 6**). The primary objective is to preserve RBC volume and ensure optimal function across the RBC’s limited lifespan. As RBCs are constantly exposed to shear stresses in circulation, the persistent activation of mechanosensitive channels may continuously lead to the disruption of the cytosolic cation composition which impacts cell volume.

**Figure. 6:**
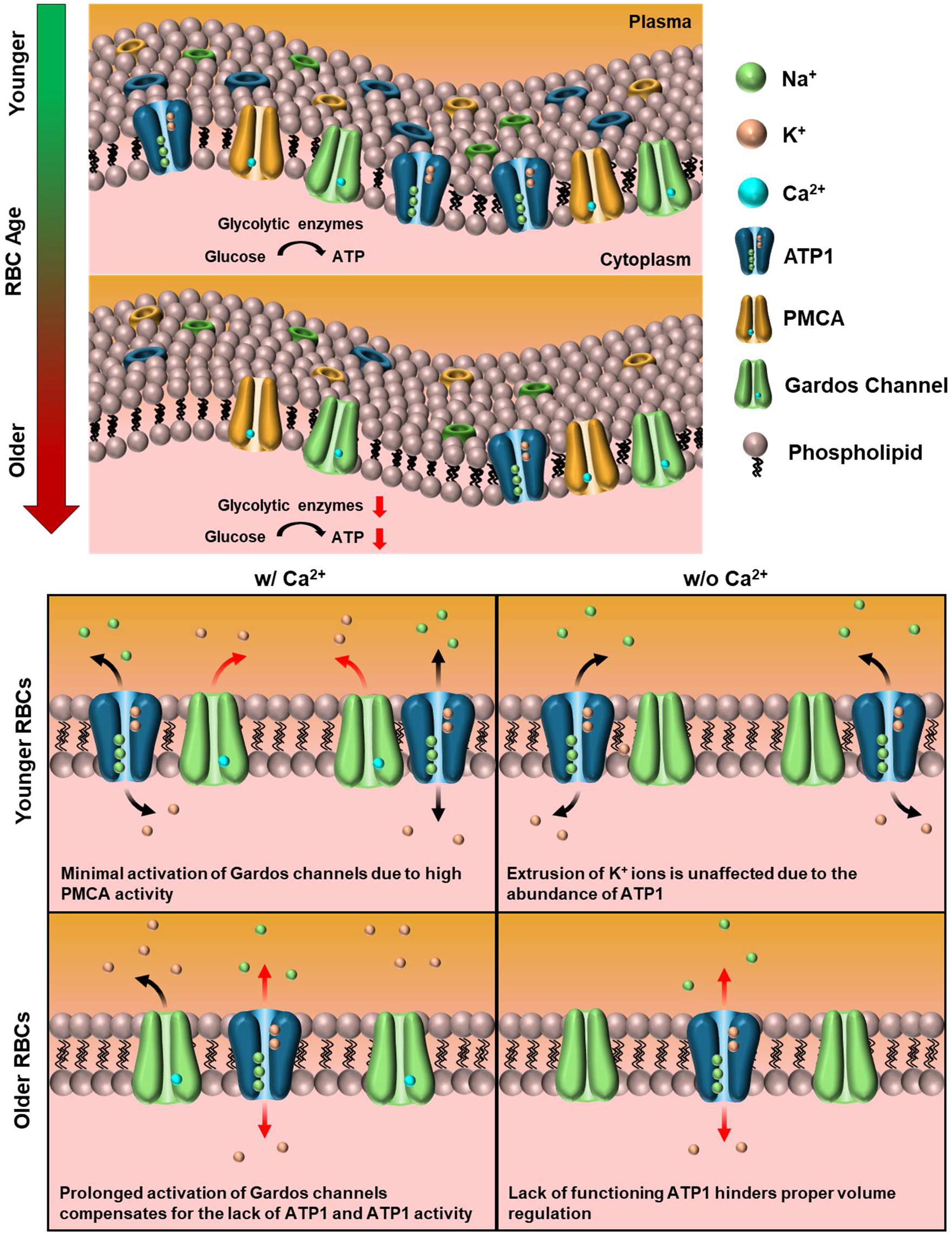
Compensation strategy for effective volume regulation in ageing RBCs. The dependence on Ca^2+^ for effective volume regulation increases with RBC age. YRBCs may not require Ca^2+^ for volume regulation due to the abundance of ATP1 and ATP. However, ORBCs with severely reduced ATP1 and ATP are unable to facilitate changes in RBC volume during fluid shear in hypocalcemia. This indicates that ORBCs dependence on Ca^2+^ is higher than YRBCs when attempting to maintain optimal shape and function.

We investigated this phenomenon by exposing RBCs to sustained shear stress to simulate *in vivo* conditions in the circulation. The difference in volume regulation between RBC subpopulations (YRBCs and ORBCs) was particularly notable when ORBCs were sheared under a hypocalcemic condition, highlighting their diminished Ca^2+^-independent pathways of volume regulation. This presents the key evidence of the disparity in volume regulation and reliance on Ca^2+^ that these two different RBC subpopulations possess. The modulation of cation fluxes across the membrane is firmly maintained by both active and passive membrane transporters such as ATP1 and Gardos channels. The cytosol of the RBC maintains a higher K^+^ concentration (133.7 vs 4.1 mM), a lower Na^+^ concentration (16.5 vs 151.6 mM)^37^ and a lower Ca^2+^ concentration (40 nM vs 1.8 mM)^17^ than the extracellular plasma. In our experiments, it is likely that the passive movement of K^+^ out of the RBC coupled with the active extrusion of Na^+^ helped RBCs to mitigate swelling caused by the excess entry of monovalent cations during shear. We confirmed that the lack of K^+^ extrusion in the case when ORBCs were sheared in the absence Ca^2+^ was likely responsible for the apparent swelling which necessitates Ca^2+^ in ORBCs.

Unlike ORBCs, YRBCs did not show any notable shear-induced size changes regardless of the suspending media. In this case, the abundance of glycolytic enzymes and ATP would allow for quick volume adjustments with the aid of ATP1. Coupled with passive movements of the cations, rapid active exchange of Na^+^ and K^+^ against their respective concentration gradients regulates the total cation levels in the cytosol^38^. Furthermore, there may be a low level of Gardos activation as the high levels of ATP-dependent PMCA activity are capable of removing excess Ca^2+^ quickly^39^. This prevents additional K^+^ in the cytosolic space from being extruded which may cause cell shrinkage. As for ORBCs, the depleted ATP levels^21^ and reduced presence of ATP1^22^ may have limited the effectiveness of ATP1 in volume regulation. Furthermore, the gradual dissipation of the transmembrane concentration gradient^40^ with age could possibly hinder the function of ATP1. Collectively, these retard the process of volume regulation through Ca^2+^-independent pathways which resulted in the swelling of the ORBCs as we observed.

The contribution of ATP to RBC volume regulation was assessed with use of MD cells. Our longitudinal study across 40 mins showed that both MD YRBCs and MD ORBCs exhibited an increased τ and an attenuated *RP* caused by reduced intracellular ATP. The notable swelling of sheared MD YRBCs due to reduced intracellular ATP supports the role of ATP-dependent transporters such as ATP1 in RBC volume regulation. It is worthy to note that MD YRBCs exhibited a significantly higher *RP* than MD ORBCs which further emphasizes that the loss of transmembrane concentration gradients due to biological ageing likely interfered with the functions of ATP1.

The retention of higher intracellular Ca^2+^ found in ORBCs after shear treatment as observed from our results corroborated several previous studies^41–44^. While this confirms a diminished PMCA function, it also suggests a sustained activation of Gardos channels which may serve to balance the excess cytosolic cations through facilitating the removal of K^+^, allowing ORBCs to regulate their volume. Our parametric study on shearing of ORBCs with different Ca^2+^ concentrations confirmed a dependence on Ca^2+^ for optimum volume regulation in ORBCs. Specifically, there appears to be a threshold (∼ 2 mM) of medium Ca^2+^ required for the activation of sufficient Gardos channels to prevent swelling of the cells. This finding aligned well within the known physiological range of Ca^2+^ (1.8 – 2.6 mM)^17,45,46^. The important involvement of Ca^2+^ in the volume regulation of ORBCs was further quantified via the inhibition of Gardos channels. As the swelling of the Gardos-inhibited ORBCs was similar to ORBCs sheared in the absence of Ca^2+^, our findings validated the contribution of Ca^2+^ and Gardos effect in regulating the volume of ORBCs. Interestingly, the activation of the Gardos channels under physiological shear stress and Ca^2+^ level did not result in dehydration as reported by other studies that involved chemical treatment of the RBCs^47–49^, suggesting an adaptive regulatory pathway to preserve the RBC size in systemic circulation.

The impact of RBC swelling on its function was also investigated based on (1) flow dynamics and (2) O_2_ delivery. For flow dynamics, swelling of ORBCs under hypocalcemic conditions resulted in a significant alteration in their movement near the bifurcation. Swelled ORBCs exhibited significantly lower *j* and *m*, underscoring the broader impact of swelling on RBC migration in circulatory flow. The relationship between RBC size and migration can be typically described using Stokes’ number, which predicts that larger particles with a higher inertia are less responsive to changes in flow direction. Consequently, they are more likely to deviate from their original streamline and migrate further away from the channel walls at bifurcating points. This principle explains why YRBCs, which are inherently larger than ORBCs, exhibited higher baseline *j* and *m* values, regardless of the suspending media. However, a Stokes number-based evaluation alone does not account for the increased *j* and *m* observed in our sheared cases given the preservation of RBC size. Moreover, it also contradicts with the decreased *j* and *m* observed in swelled ORBCs when sheared in the absence of Ca^2+^, suggesting that intrinsic cellular properties, particularly RBC deformability may play a more critical role in determining the cell’s trajectory.

Deformability influences the wall lift force^50^ and centreline flow migration rate^51^ of RBCs which was quantified in our study by *j* and *m*, respectively. Combining the effects of accumulated oxidative damage^52^ and increased cytoplasmic tortuosity^53^, ORBCs are expected to be less deformable than YRBCs which is consistent with our baseline DI measurements (**Supplementary Table S2**). It is likely that YRBCs exhibited a more pronounced tank-treading motion due to its higher deformability that facilitated the ability to cross flow streamlines, propelling them further away from the vessel walls and increasing their radial migration rate toward the flow centreline^54^. In contrast, ORBCs, with their lower deformability, would experience a reduced frequency of tank-treading^55^, resulting in a lower initial displacement away from the channel wall post-bifurcation and reduced inward migration rate toward the flow centreline downstream as compared to YRBCs. The increased deformability after shear treatment may have facilitated a higher frequency of tank treading motion which led to the increase in *j* and *m*. As deformability of RBC can also be affected by changes in the intracellular viscosity and hydration state of the cell^56^, swelling of ORBCs likely contributed to the deterioration of their biophysical properties, outweighing the shear-induced increase in deformability.

RBC O_2_ affinity and O_2_ release rate were investigated to determine the impact of swelling on O_2_ delivery capacity. The p50 of YRBCs was significantly higher than that of ORBCs, akin to what was reported previously^57^. Our p50 findings suggest that the swelling phenomenon did not have bearing on the p50 of ORBCs as all the cases revealed a reduction in p50 after shear treatment, indicating an increased affinity of hemoglobin to O_2_. On the other hand, ORBCs was observed to have a faster O_2_ release rate than YRBCs, consistent with previous findings^58^. The reduced MCV in ORBCs^59^ contributes to a shorter cytoplasmic pathlength, thereby lowering the time that is required for O_2_ to diffuse out of the cells. Swelled ORBCs revealed a slower O_2_ release rate, which confirmed that cytoplasmic pathlength may be playing a dominant role in controlling the O_2_ exchange of RBCs^60^.

In conclusion, our study revealed a fundamental shift in RBC volume regulation with ageing. While YRBCs rely on ATP-dependent mechanisms to maintain volume under shear stress, ORBCs become increasingly dependent on Ca²⁺-mediated pathways, reflecting an adaptive yet vulnerable response to declining metabolic capacity. This shift has direct consequences on microcirculatory function. Under hypocalcemic conditions, failure of Ca²⁺-dependent regulation leads to persistent swelling in older RBCs, impairing deformability, increasing flow resistance, and altering their distribution in the microvasculature. Concurrently, swollen RBCs exhibit slower oxygen release, reducing oxygen availability to endothelial cells. Together, these findings establish a link between RBC ageing, ion regulation, and hemodynamic function, suggesting that disturbances in Ca²⁺ homeostasis may contribute to microcirculatory impairment and potentially vascular disease.

## Materials and Methods

### Blood collection and sample preparation

This study was approved by the National University of Singapore Institutional Review Board (NUS-IRB-2022-114) and all procedures were performed in accordance with the approved guidelines. Healthy male volunteers ranging from 21 to 30 years of age, without any known pathological conditions that could affect the properties of the collected blood, were enlisted for this study. Informed consent was obtained from all recruited participants. All blood collection and storage processes were performed under sterile conditions. Venous blood (20 mL) was collected into ethylenediaminetetraacetic acid coated tubes.

RBCs were separated from whole blood by centrifugation at 2500 x *g* for 10 min (Megafuge 8, Thermo Scientific, Massachusetts, USA). The isolated RBCs were washed thrice with 1X phosphate-buffered saline (PBS). To assess the effects of *in vivo* ageing, the washed RBCs were further density-fractionated at 4600 x *g* for 30 min to obtain the younger RBCs (top 5% fraction) and older RBCs (bottom 5% fraction)^58,61^. To assess the contributions of Gardos effect in volume regulation, the respective subpopulations of RBCs were treated in PBS with Senicapoc (10 µM) (SML2500, Sigma Aldrich, Missouri, US) for 10 min^62^.

### Couette flow shearing

RBCs were suspended in PBS at a hematocrit of ∼1%. A cone-and-plate rheometer (DVII+ Pro, Brookfield, Toronto, CA) was used to generate a couette flow and impose a constant shear stress distribution throughout the test samples. Shearing of the samples was performed under the shear condition of 15 Pa for 300 s at 37°C. This shear stress condition was selected for exposing the RBCs to the upper limit of physiological shear stresses found in the cardiovascular system.

### Machine learning-based image analysis

In order to obtain an accurate and fast RBC size determination, we employed a 2-dimensional size quantification approach from microscope images, processed with a machine learning-aided image segmentation algorithm. RBCs were imaged using an inverted microscope (IX-71, Olympus, Tokyo, JP) at a magnification of 96X. For determination of RBC size, RBC images were taken using an 8-bit monochrome CMOS camera (FASTCAM-1024 PCI, Photron, Tokyo, JP). At least 300 individual RBCs were imaged for each sample from which the population distribution statistics were derived. A deep learning convolutional neural network (CNN) model was used for image classification to accurately identify RBC discocytes for size analysis. CNN models including residual net (ResNet50), SqueezeNet, EfficientNet, GoogleNet, and visual geometry group (VGG19)^63,64^ were compared to select the most appropriate model for our purpose. A total of 1,440 single RBC images (480 images per each cell type) were utilized for this selection process. 70% of the dataset were used for training while the rest were used for validation purposes. Artificial augmentation operations such as random flipping and rotations were performed on the training dataset to improve the robustness of the CNN models and to reduce the potential issue of overfitting. A total of 3 epochs were carried out for each model using the same training dataset with the dataset being split into 14 even and random iterations for training prior to validation. For the testing phase, three separate sets consisting of 500 single RBC images were prepared. Each image in the test sets was manually labelled by two trained individuals to assess the accuracy of our CNN models. A binary classification confusion matrix for each test set was calculated to evaluate the performance of each CNN model to determine the overall accuracy. Automated image segmentation and blob analysis was then used to extract the area information of labelled discocytes (**Supplementary Figure. S1**). The use of two-dimensional area to represent RBC size was validated by comparison with MCV measured using an automated hematology analyzer (XN-3100, Sysmex, Kobe, Japan).

We showed that our method was comparable to the golden standard of MCV measurements used in hospitals, affirming the reliability of our results. Interestingly, the strong linear correlation between the area and MCV measurements corresponded with a gradient of ∼2 µm which is akin to the known thickness of an RBC^65^ (**Supplementary Figure. S2**). The method’s robustness was further assessed by evaluating the reproducibility of area measurements for the same RBC samples at two different time points (t = 0 and 40 mins) (**Supplementary Figure. S5**). This also confirmed that prolonged *ex vivo* incubation at 37°C did not affect RBC size.

### Quantification of size recovery dynamics

The recovery dynamics (**Figure. 2b**) of swelled RBCs were quantified through two parameters derived from a fitted exponential curve (*P* = Ae^−βt^ + α): (1) recovery rate constant, 0 and (2) recovery potential, 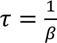 and (2) recovery potential, 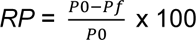, where *P*0 represents the percentage size change of the cell immediately after shearing and *Pf* is the percentage size change 40 min after shearing. The rate constant for the recovery of RBC size after shear treatment was quantified by τ, where a higher τ indicated a slower recovery. The extent to which the RBCs were able to return to their original size after shear treatment was defined as *RP*.

### Measurement of extracellular Ca^2+^ and K^+^ concentration

Medium Ca^2+^ and K^+^ concentrations were quantified using a blood gas analysis system (epoc, Siemens, Munich, DE). Different Ca^2+^ concentrations (0.5, 1.0, 1.5 and 2.0 mM) were prepared by dissolving CaCl_2_ in PBS prior to analysis. The concentration of K^+^ ions in the supernatant of the sample was measured before and after shear treatment to determine the efflux of K^+^ from the sheared RBCs.

### Measurement of intracellular Ca2+ concentration

Intracellular Ca2+ level was determined with a Ca2+ indicator (Fluo4-AM, Thermo Fisher Scientific, Waltham, USA). Washed RBCs were incubated in Fluo4-AM (5 µM) for 60 min at 37°C, protected from light. Fluo4-loaded RBCs were washed thrice with PBS before being suspended in PBS with Ca2+ at a hematocrit of ∼1%. The RBC suspension was placed in the dark at 37°C for 20 min to allow for the intracellular esterases to cleave and activate the dye. The loaded RBCs were then excited at λ = 488 nm. The emitted fluorescence signal was detected at λ = 505 nm and observed using confocal microscopy (FV1000, Olympus, Tokyo, JP) at 100X magnification. Quantitative measurements of Fluo4 fluorescence were performed using a multimode microplate reader (Varioskan™ LUX, Thermo Fisher Scientific, Massachusetts, USA). Fluo-4 fluorescence before and after shearing were compared to determine the degree of Ca2+ retention in each subpopulation of RBCs after shear treatment.

### Measurement of intracellular ATP

ATP levels were measured with a luminescence-based ATP assay kit (A22066, Thermo Fisher Scientific, MA, USA). The RBC samples were assayed at two time points: (1) immediately after density fractionation and (2) after 8 hr of incubation in a 37°C water bath (MD cell preparation). The absorbance was determined using a multimode microplate reader (Spark ®, Tecan Group Ltd., Männedorf, CH).

### RBC deformability

Ensemble-averaged RBC deformability was determined with Ektacytometry (Rheoscan AnD, RheoMediTech, Seoul, KR). The deformability index (DI) was used to quantify the deformability of RBCs.

### Single cell dynamics across in vitro bifurcation

To assess the effects of shear-induced volume change on single RBC flow dynamics, an *in vitro* microchannel mimicking an *in vivo* arteriolar bifurcation was fabricated using polydimethylsiloxane (PDMS, Dow Corning, MI, USA) by standard soft lithography and replica moulding techniques (**Supplementary Figure. S3**). The respective subpopulations of RBCs were evaluated before and after shearing, in media with and without Ca^2+^. The upstream lateral positions of the RBCs in each experiment were first initialized towards the inner wall of the main channel (**Supplementary Figure. S4**) by a sheath fluid containing the same suspending media, introduced from a separate inlet. The baseline 0D was determined at the end of the filleted corner and kept constant across each experiment. The inlet flow rates for the RBCs and sheath fluid were maintained with volumetric syringe pumps (KDS210, KD Scientific, Massachusetts, USA) at 0.2 and 3.4 µL/min, respectively to match a pseudo shear rate of 200 s^-1^. The shear stress distribution across the microchannel was estimated from two-dimensional computational fluid dynamics simulations using a commercial software (COMSOL Multiphysics 5.1, Stockholm, SE). A viscosity of 1.01 cP and density of 999.6 kg/m^3^ were used to define the fluid domain rheology to match our experimental conditions. Mesh dependence testing was performed to optimize the mesh density for estimation of the shear stress experienced by RBCs during our experiment (**Supplementary Figure. S6**).

### Measurement of p50 and O_2_ diffusion

p50 was quantified via measurement of dissolved gases and media pH using a blood gas system (RAPIDPoint 500e, Siemens, Munich, DE). The O_2_ release rate from RBCs was measured using a custom-designed set-up (**Figure. 5c**) containing an O_2_ microsensor (OX-10, Unisense, Aarhus, DK). The O_2_ sensor was used to measure and monitor the concentration of dissolved O_2_ in the RBC suspension. The sample volume was kept constant for all cases with a fixed hematocrit (∼1%). Prior to the start of the experiment, each sample was initialized with sterile air for 15 min to ensure that the sample solution has reached an O_2_ saturation of 283 µmol/L. To initiate the RBC deoxygenation, nitrogen gas (N_2_) was then bubbled at a constant pressure of 200 mbar with the use of a pressure controller (OB1, Elveflow, Paris, FR). The concentration of dissolved O_2_ was recorded at 20s intervals across a fixed duration (300s) to determine the rate of deoxygenation in each sample. Quantification of the RBC O_2_ release rate was derived from the fitted exponential fit (*_C_* = 283*_e_*^−αt^) where α represents the decay rate of the exponential fit (O_2_ release rate). A higher α represents a faster O_2_ release rate. Temperature was maintained at 37°C throughout the experiments.

### Statistical analysis

Statistical analyses were performed using a commercially available data analysis software (OriginPro 2022, OriginLab, MA, USA). p < 0.05 was considered to be statistically significant. All physiological and rheological values are presented as means ± SEM. The Kolmogorov-Smirnov test was used for normality test. Paired and unpaired student’s t-tests were conducted for two-group comparison. Two-way ANOVA or two-way repeated measures ANOVA with post-hoc Tukey’s comparisons was applied for comparison of more than two groups. Slope significance testing was performed using analysis of covariance (ANCOVA).

## Acknowledgments

JL was supported through the NUS Graduate School Integrative Sciences and Engineering Program. The MCV measurements was assisted by the Singapore General Hospital. **Figures. 5c and 6** were produced with the aid of Biorender. This research was supported by the Singapore Ministry of Education MOE) University Research Committee Tier 2 grant MOE-T2EP50121-0015.

